# The Mismatch Between Neuroscience Graduate Training and Professional Skill Sets

**DOI:** 10.1101/2021.08.09.455678

**Authors:** S. Shah, A.L. Juavinett

**Author notes:** **Corresponding author:** Ashley L. Juavinett, PhD, York Hall 4070C, Mail Code #0355, UC San Diego, 9500 Gilman Drive, La Jolla, CA 92093.

## Abstract

Understanding the skill sets required for career paths is a prerequisite for preparing students for those careers. Neuroscience career paths are rapidly changing as the field expands and increasingly overlaps with computational and data-heavy job sectors. With the steady growth in neuroscience trainees and the diversification of jobs for those trainees, it is important to assess whether or not our training is matching the skill sets required in the workforce. Here, we surveyed hundreds of neuroscience professionals and graduate students to assess their use and valuation of a range of skills, from bench skills to communication and management. We find that professionals with neuroscience degrees can be clustered into three main groups based on their skill sets: academic research, industry research and technical work, and non-research. Further, we find that while graduate students do not use or highly value management and communication skills, almost all neuroscience professionals report strongly needing those skills. Finally, coding and data analysis skills are widely used in academic and industry research and predict higher salaries. Our findings can help trainees assess their own skill sets as well as encourage educational leaders to offer training in management and communication–skills which may help catapult trainees into the next stages of their careers.

## Introduction

The past two decades of neuroscience research are marked by dramatic innovations in our ability to record, manipulate, and predict brain activity (Luo et al., 2018; Sejnowski et al., 2014). In addition to expanding the size of our datasets, these innovations are also changing what kind of therapeutics are being developed, how artificial intelligence is implemented, and what kind of evidence is permissible in a courtroom. As a result, there is an increasing need for people in the workforce who understand the process and outputs of neuroscience research and who can communicate this to public shareholders. In parallel, more and more students are graduating with degrees in neuroscience (Rochon et al., 2019). These degree holders are finding themselves in a variety of roles and job sectors, including applied and industry research, policy making, and consulting, reflecting the growing needs of our society.

Harkening back to the early days of artificial intelligence, neuroscience is also deeply integrated with computer science and the rapidly growing fields of data science and machine learning (Paninski & Cunningham, 2018; Akil et al., 2011). This trend, combined with the increasing size of neuroscience datasets, has inspired many leaders in neuroscience education to call for an increase in quantitative skill sets, particularly coding (Ramirez, 2020; Akil et al., 2016; Grisham, 2016). Beyond neuroscience, technical skills such as coding are becoming essential in our increasingly automated economy (Cummins et al., 2019).

As more students earn undergraduate and graduate degrees in neuroscience, it is important that we take into consideration the possible career paths and requisite skill sets for these graduates (Rochon et al., 2019). Gaps between the training and the demands of a profession can result in a lack of job preparedness as trainees enter the workforce (Suleman, 2017; Wendler et al., 2012; Bridgstock, 2009). In 2009, the Springboard Project American Workforce Survey revealed that half of the adults surveyed reported a gap between professional needs and skills of employees; additionally, the surveyors described a deficit in “soft skills” like work ethic, communication, and accountability (Business Roundtable, 2009). In STEM fields specifically, skills such as communication, project management, teamwork, problem solving, critical thinking, and interpersonal skills have been repeatedly reported as lacking in trainees (Hung-Lian et al. 2000; Radermacher and Walia 2013). Skill assessments can help educators identify possible gaps within different fields by shedding light on the expectations and needs of the workforce, and could be especially useful in quickly evolving fields such as neuroscience.

For example, a previous study by Cui & Harshman (2020) performed a skill assessment on chemists in different job sectors to determine which skills are required to succeed in their respective professions. Interviewing chemists in the academia, industry, and government sectors, Cui & Harshman (2020) grouped knowledge and skills into twelve main themes and noted the importance of skills such as communication and management regardless of job sector. The authors conclude that while certain skills are rightfully emphasized in the training of chemists, it would also be beneficial to provide more catered skill training based on trainees’ intended career paths.

Similarly, the diverse and ever-evolving nature of neuroscience career paths requires educators to ask if neuroscience education and training is adequately preparing students with both quantitative and soft skill sets to satisfy the needs of their future job sector. To address this question, we assessed the skills of professionals with neuroscience degrees to understand the mastery, frequency, and importance of various technical and so-called “soft skills” in their respective fields. This analysis highlights the importance of computational, analytical, managerial, and communication skills and can inform educational leaders and mentors on how to provide more efficient training for future neuroscientists.

## Materials & Methods

### Participant recruitment

A 50-question survey was administered online via Qualtrics and distributed via social media and email. Participants either held at least one degree (B.S., B.A., M.S., or Ph.D.) in neuroscience or related fields (e.g. Cognitive Science) and/or 1+ year experience conducting neuroscience research, defined as “any research pertaining to the nervous system.” While the scope of this survey encompassed neuroscientists at various educational and career levels, we limit our analysis and discussion here to responses from current graduate students as well as neuroscience professionals (e.g., faculty members, industry professionals) to understand which skill sets are being utilized in these groups.

As shown in Table 1, we included responses from 116 current graduate students as well as 125 neuroscience professionals, including 57 faculty members and 68 participants in a wide range of other fields including government, industry research, and science communication. Within the graduate student participants, 65 were current PhD students, 5 were current Master’s students, and 46 did not specify. “Faculty member” includes research faculty only; “teaching faculty” or “instructors” are included within “Science Communication & Teaching.” Participants were asked to self-describe their current position into provided job titles. Those that answered “Other” were sorted according to their position title. Several related job titles were grouped, such as “Data Science” and “Software Engineer” due to low numbers of responses in individual categories. Our final grouping includes eight different job categories (Table 1).

**Table 1.** Study participants self-identified their Job Sector (first column). The number of participants per sector is shown in the N_sector_ column. Representative job titles are also provided to illustrate the types of positions represented in our dataset.

| Job Category | Representative Position Titles | N<sub>sector</sub> |
| --- | --- | --- |
| Applied/Industry Scientist | Scientist (Research, Principal); User Experience Researcher | 13 |
| Consultant | Scientific Consultant; Senior Analyst | 7 |
| Data Science & Software Engineering | Data Scientist; Analyst (Strategy, Senior); Software Engineer | 13 |
| Faculty Member | Professor (Assistant, Associate, Full); Assistant Investigator | 57 |
| Graduate Student | Masters student; PhD student | 116 |
| Industry Sales & Marketing | Sales Manager; Account Manager | 3 |
| Management | Project Director; Scientific Program Manager; Health Scientist Administrator; Director of Innovation Policy | 17 |
| Medical Professional | Psychologist (Clinical, Neuro); Assistant Professor (Clinical) | 3 |
| Science Communication & Teaching | Writer (Science, Medical); Editor (Senior, Deputy, Executive Story); Lecturer; Freelance Science Communicator | 13 |

### Survey

The entire survey assessed a wide range of demographics, background, skills, and career path information. Here, we focus on a set of questions regarding the skills necessary for these professionals in their respective fields, akin to work in previous studies (Jang, 2015). Note that this study assesses skills, rather than underlying abilities that may enable those skills. The list of skill items was developed *de novo* to capture the breadth of skills required in neuroscience research and career paths (see Table 3 for all items). In a Likert-style scale, participants were asked to describe their use of these 21 different skills. For each skill, participants were asked:

**Table 2.** Top skills for neuroscience graduate students (n=116) and professionals (n=125). Mean and standard deviation are shown in parentheses.

|  | Mastery | Frequency | Importance |
| --- | --- | --- | --- |
| **Graduate Students**<br>(n=116) | 1. Designing and planning experiments ($2.36 \pm 0.61$)<br>2. Synthesizing existing research ($2.31 \pm 0.65$)<br>3. Verbally presenting information in front of an audience ($2.28 \pm 0.72$) | 1. Communicating with other scientists and/or clients ($2.97 \pm 1.11$)<br>2. Synthesizing existing research ($2.79 \pm 1.00$)<br>3. Running statistical analyses ($2.53 \pm 0.92$) | 1. Designing and planning experiments ($3.45 \pm 0.85$)<br>2. Running statistical analyses ($3.33 \pm 0.79$)<br>3. Writing (for a scientific audience, includes writing grants and papers) ($3.25 \pm 0.84$) |
| **Professionals**<br>(n=125) | 1. Communicating with other scientists and/or clients ($2.61 \pm 0.28$)<br>2. Verbally presenting information in front of an audience ($2.35 \pm 0.32$)<br>3. Synthesizing existing research (eg. reading papers, producing literature reviews) ($2.19 \pm 0.37$) | 1. Communicating with other scientists and/or clients ($3.27 \pm 0.41$)<br>2. Managing a team ($2.70 \pm 0.46$)<br>3. Synthesizing existing research (eg. reading papers, producing literature reviews) ($2.36 \pm 0.32$) | 1. Communicating with other scientists and/or clients ($3.21 \pm 0.41$)<br>2. Verbally presenting information in front of an audience ($2.85 \pm 0.52$)<br>3. Managing a team ($2.66 \pm 0.51$) |

**Table 3.** Participants were asked to rate their mastery in as well as the importance and frequency of various individual skills. These skills were clustered into Skill Categories for analysis.

| Skill Category | Individual Skills | Mastery $\alpha$ | Often $\alpha$ | Importance $\alpha$ |
| --- | --- | --- | --- | --- |
| Bench | Molecular Biology | 0.853696 | 0.869509 | 0.869404 |
|  | Image processing and/or microscopy | 0.845626 | 0.849925 | 0.853429 |
|  | Managing lab resources | 0.857589 | 0.848231 | 0.856032 |
|  | Building or manipulating hardware | 0.846364 | 0.840746 | 0.851486 |
|  | Designing and planning experiments | 0.851135 | 0.841128 | 0.852399 |
|  | Performing surgeries on non-human animals | 0.854401 | 0.844642 | 0.853572 |
|  | Running physiology and/or behavioral experiments | 0.854401 | 0.844642 | 0.853572 |
| Coding | Coding to analyze data or run experiments | 0.897576 | 0.841235 | 0.841779 |
|  | Computational modeling | 0.885371 | 0.807250 | 0.815021 |
|  | Writing front-end code or developing software | 0.885371 | 0.807250 | 0.815021 |
| Data Analysis & Visualization | Running statistical analyses | 0.769681 | 0.786061 | 0.835464 |
|  | Generating figures | 0.769681 | 0.786061 | 0.835464 |
| Management & Communication | Managing a team | 0.678092 | 0.392719 | 0.572120 |
|  | Writing (for a non-scientific audience) | 0.628697 | 0.441684 | 0.396392 |
|  | Communicating with other scientists and/or clients | 0.621399 | 0.425576 | 0.433642 |
|  | Verbally presenting information in front of an audience | 0.621399 | 0.425576 | 0.433642 |
| Mentorship & Teaching | Mentoring students | 0.900136 | 0.827227 | 0.884174 |
|  | Teaching in a classroom setting | 0.900136 | 0.827227 | 0.884174 |
| Scientific Writing & Synthesis | Writing (for a scientific audience, includes writing grants and papers) | 0.669985 | 0.70185 | 0.714812 |
|  | Synthesizing existing research | 0.669985 | 0.70185 | 0.714812 |

- What level of **mastery** in this skill do you need to do your current job? (=None, 1=Basic, 2=Intermediate, 3=Expert)
- How **often** do you use this skill in your current job? (0=Never, 1=A few times a year, 2=Several times a month, 3=Weekly, 4=Daily)
- How **important** is this skill in your current job? (0=Not at all important, 1=A little important, 2=Somewhat important, 3=Very important, 4=Extremely important)

For numerical analysis, responses were converted to values 0 to 3 for mastery and 0 to 4 for frequency (how often) and importance.

In addition, we analyzed information about participants’ yearly salaries. Responses were dropped for any uninterpretable inputs for this question — for example, if it was not clear that the response was a salary for a given year. If respondents gave responses that were by the month or for a 9 month salary, those responses were converted to yearly salary amounts. If respondents added any caveats about health insurance or tuition, those caveats were removed and unadjusted numbers were used. If participants gave a range, the middle of that range was used. Many responses to this question were not in US Dollars (USD). Responses were converted to USD according to exchange rates in October 2020. In order to identify outlying salaries, all salaries were Z-scored. Salaries with a Z-score greater than 2 were removed, resulting in the removal of 6 outliers.

### Statistical Approaches

To reduce the dimensionality of our dataset and determine appropriate groupings for the skill sets of professionals in different job sectors, we first Z-scored the data such that it had a mean of 0 and a standard deviation of 1. We then ran PCA (# components = 3) and K-means analyses using standard Scikit-Learn packages in Python. To determine differences between skill mastery, frequency, and importance between graduate students and three job sectors, we ran Dwass-Steel-Critchlow-Fligner pairwise comparisons. Given that we tested for six different skills for each question, we used a Bonferroni correction to determine an appropriate alpha value (0.05/6 comparisons). We therefore considered pairwise comparisons significant with a p-value less than 0.008. To identify relationships between salary and skills in specific domains, a Pearson correlation value was calculated. Relationships were considered significant with p-values less than 0.05.

## Results

First, we computed averages for mastery, frequency, and importance across our 21 different skill sets. On average, graduate students highly ranked various skills, most notably ‘Designing and planning experiments,’ ‘Synthesizing existing research,’ and ‘Running statistical analyses.’ The top skills were very different in professionals with neuroscience degrees. Across all professionals, ‘Communicating with other scientists and/or clients’ was the highest ranked skill in mastery, frequency, and importance. ‘Verbally presenting information in front of an audience’ was the second highest skill in mastery and importance for all professionals, while ‘Managing a team’ was the most often used.

There was notable variability across both graduate students and professionals, likely reflecting the variability in graduate school expectations and perceptions, as well as the differences in skills used in different career paths. To look for patterns in different career paths, we compared the skill profiles of graduate students along with eight different job categories (Figure 1). Notably, the skills required in graduate students are visibly broad and different from those required by all neuroscience professionals. Graduate students report using and needing mastery in almost all of the bench skills we assessed, whereas within neuroscience professionals, only faculty members report using and needing these skills. Faculty also reported high mastery, frequency, and importance across a wide variety of skills, with higher reported averages in almost every category than professionals in other job sectors. Coding, computational modeling, and developing software were used in several different career paths, including applied/industry science and consulting.

**Figure 1.**
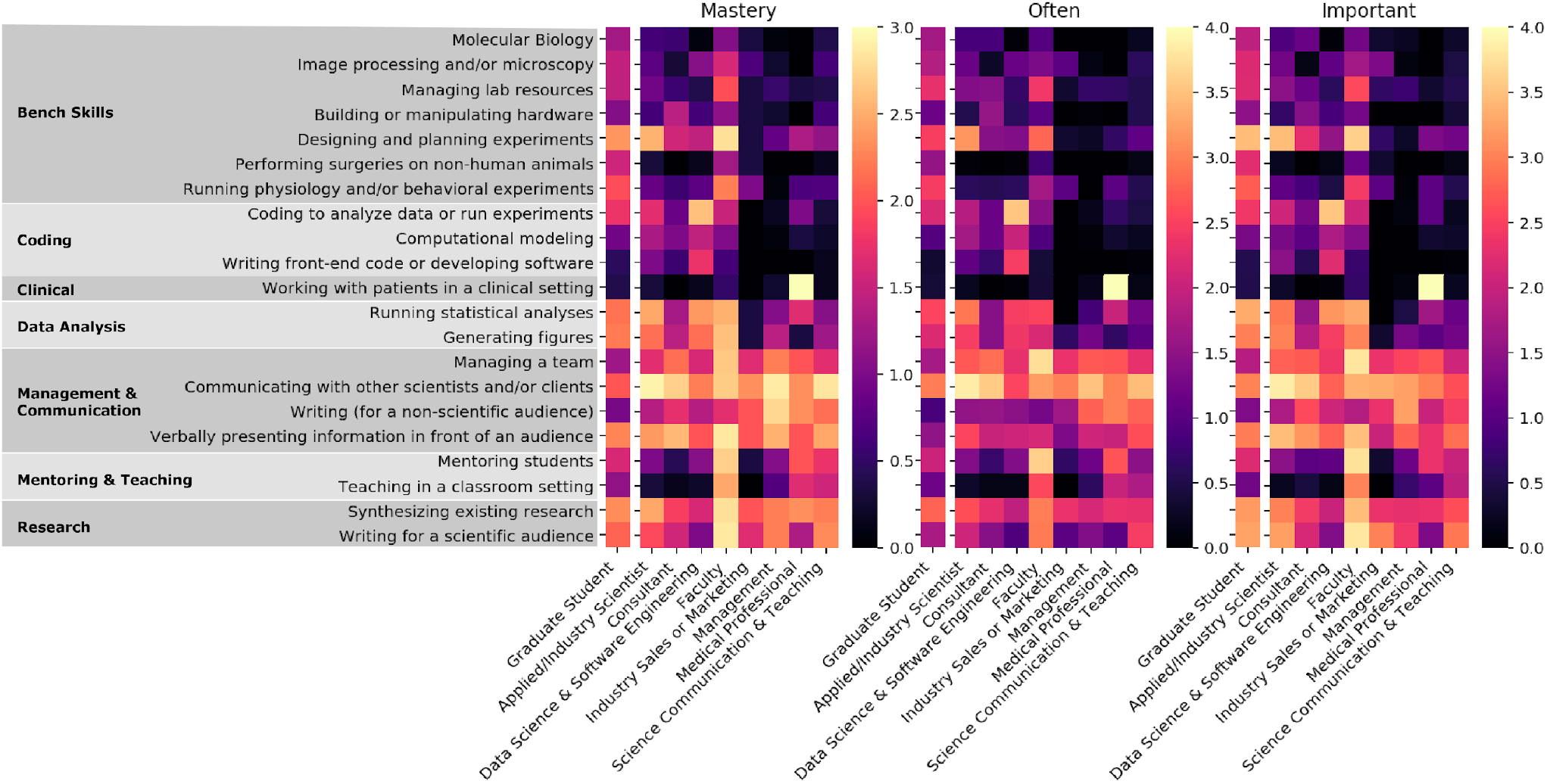
Mastery, frequency, and importance of 21 different skills across graduate students and different job sectors. Graduate students can be seen in the first, separated column. Skills are ordered based on their final groupings. Survey responses were converted to numbers for analysis Mastery: 0=None, 1=Basic, 2=Intermediate, 3=Expert. Frequency: 0=Never, 1=A few times a year, 2=Several times a month, 3=Weekly, 4=Daily. Importance: 0=Not at all important, 1=A little important, 2=Somewhat important, 3=Very important, 4=Extremely important.

### Twenty-one skill items can be reduced to fewer skill categories

Next, we sought to reduce the dimensionality of our 21-skill profiles for further analysis. We computed a cross-correlation for all of the skills for all participants for each question to determine whether some skills were highly correlated with each other (Importance shown in Figure 2). This analysis, along with the conceptual relationships between these skills, suggested that we could reduce our 21 skill items to fewer skill categories. Although participants were also asked about ‘Working with patients in a clinical setting,’ only Medical Professionals gave this category scores higher than 0 and these responses did not correlate with any other skill. It was therefore excluded in subsequent analyses, resulting in six final skill categories: bench, coding, data analysis, management and communication, mentorship and teaching, and research (Table 3).

**Figure 2.**
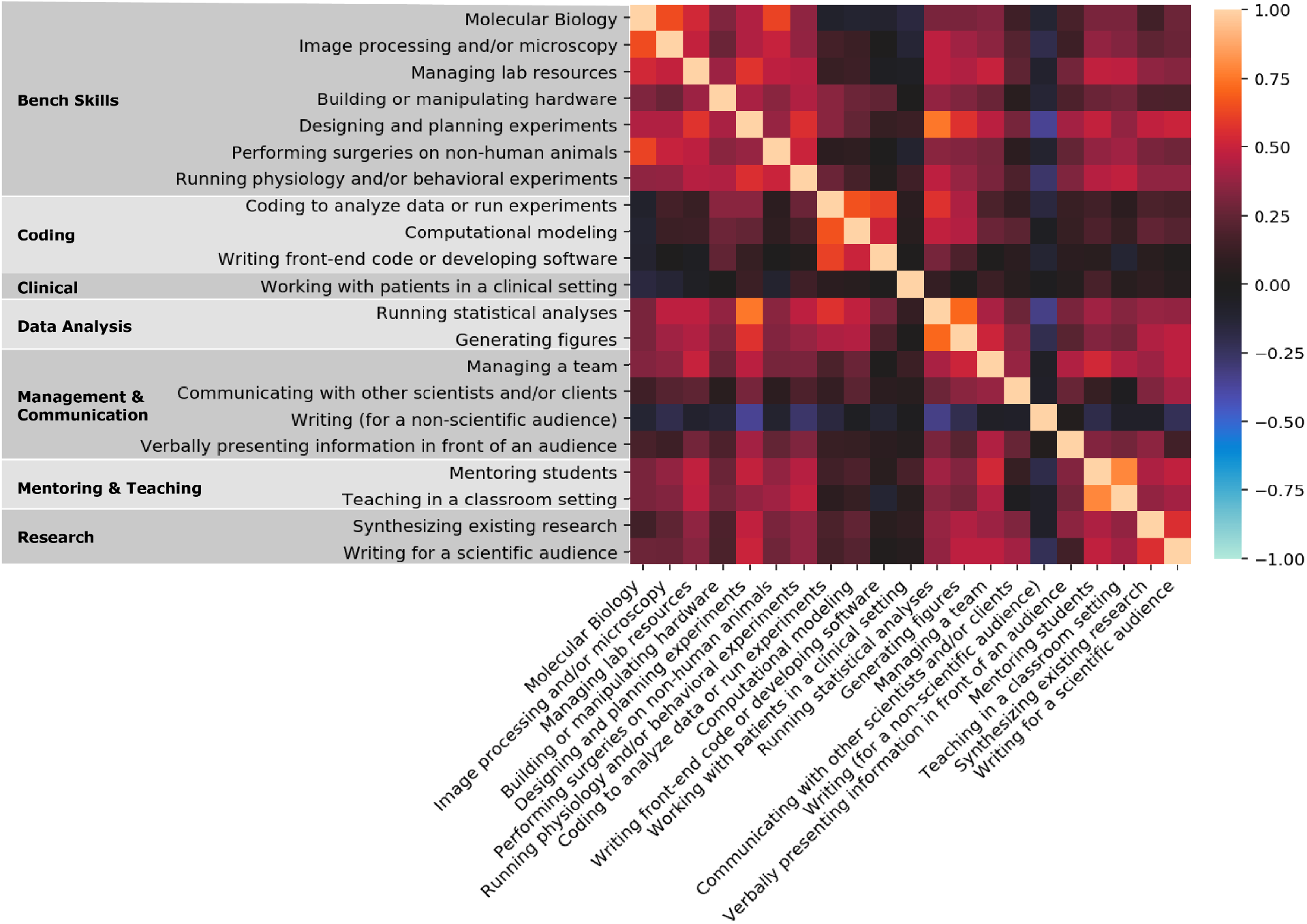
Correlation heatmap for the importance ratings of all 21 skills for all professionals. Skills are ordered based on their final groupings. Mastery and frequency heatmaps were almost identical.

To confirm the statistical robustness of our skill category groupings, we computed a Cronbach’s alpha for each category, for each of the three questions. Doing so resulted in a Cronbach’s alpha higher than 0.60 for each category, with the exception of the frequency and importance of management & communication skills (Table 3). This likely reflects the diversity of skills that are included in this category.

### Individual skill profiles can be clustered based on job sector

In any effort to understand if individual skill profiles could be clustered in an unbiased way, we then used a PCA to visualize participant’s skill profiles in fewer dimensions (Figure 3). Based on the categorization of each individual’s reported job sectors into the eight job sectors, the PCA plots for mastery, frequency, and importance show that most job sectors are discernibly clustered and that the job sectors occupy different skill spaces. A K-Means clustering analysis confirmed that we could reduce these job sectors into three large groups:

**Figure 3.**
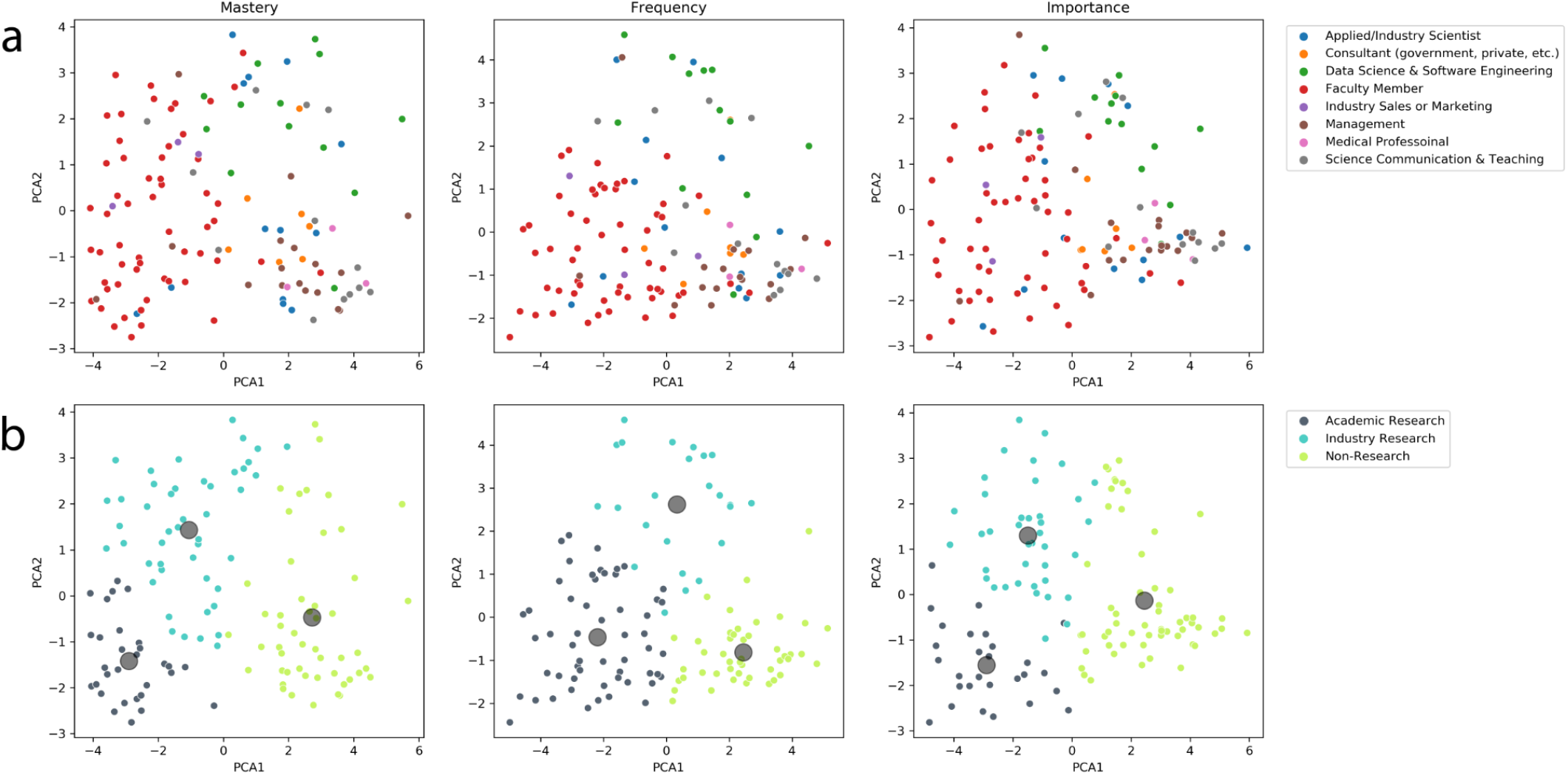
Reducing the dimensionality of skill sets into larger job sectors. **a**. First two PCA dimensions after analyzing all participant skill profiles. **b**. K-means clustering of PCA projection, demonstrating that skill profiles can be loosely clustered into three separate groups: academic research, industry research, and non-research.

- **Academic Research:** Faculty Member
- **Industry Research:** Applied/Industry Scientist, Data Science & Software Engineering
- Non-Research: Consultant, Management, Science Communication & Teaching, Industry Sales or Marketing, Medical Professional.

These clusters were most clearly distinguished in how often these skills were used (middle column, Figure 3b) and reflect the differences in skill profiles as highlighted in Figure 1.

### Mastery, frequency, and importance of skills varies across job sectors

Once we were confident that the skill profiles could be clustered into job sectors, the averages of the mastery, frequency, and importance rankings of each skill set were plotted for graduate students as well as each of the job sectors (Figure 4). Several interesting findings emerged from this analysis.

**Figure 4.**
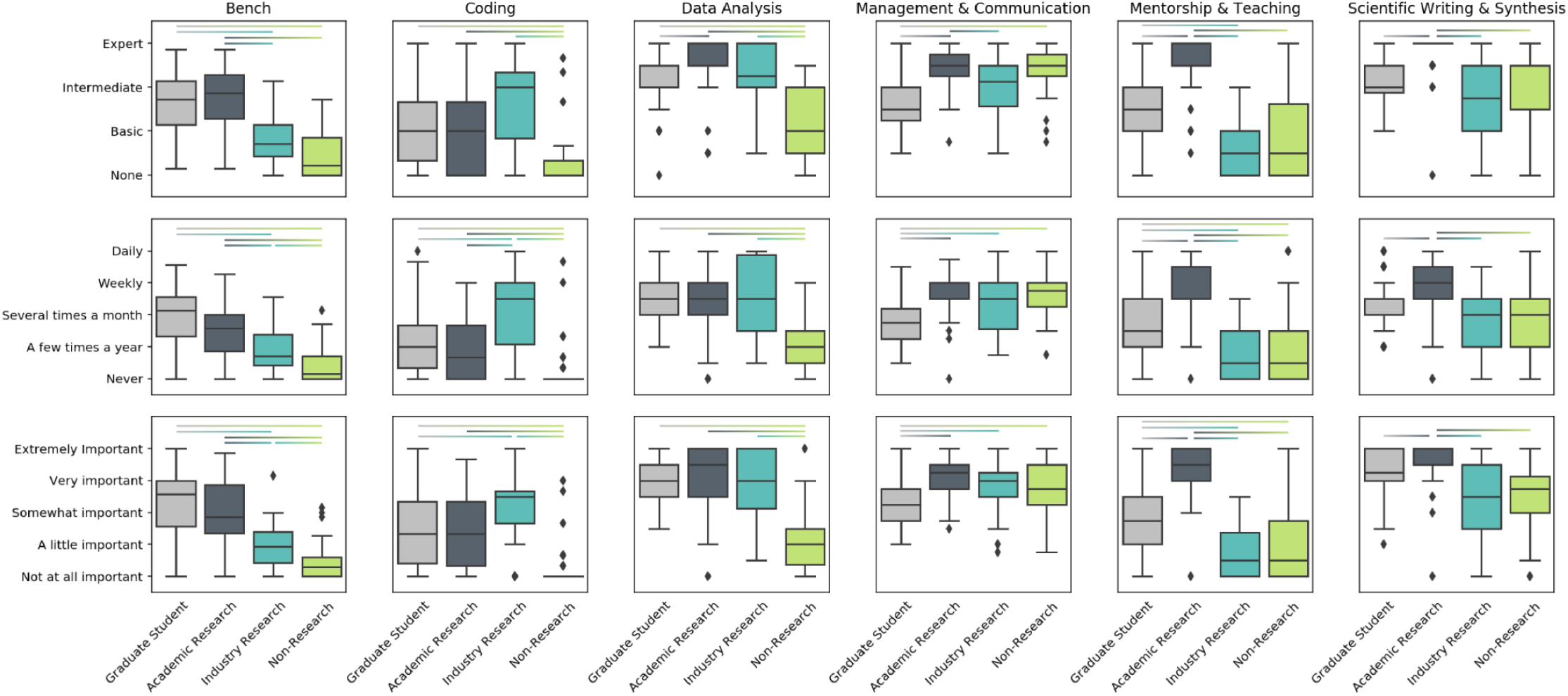
Graduate students and three different job sectors value and practice different skill sets. Median mastery (top row), frequency (middle row), and importance (bottom row) for six skill sets across graduate students (light gray) and three different job sectors (academic research: dark gray; industry research: teal; non-research: lime). Lines above box plots indicate p<0.008 as tested with a Dwass-Steel-Critchlow-Fligner test (see Methods for Details)

While Bench skills were frequently used, important, and required by graduate students and faculty, they were less prevalent in Industry research and Non-Research sectors. All sectors besides Non-Research jobs reported needing at least a basic understanding of Coding, and noted that coding was a little bit important. This was highly variable even among researchers though, likely reflecting the fact that not all research requires coding experience. All sectors except Non-Research rated Data Analysis and Scientific Writing & Synthesis skills somewhat highly, though these did show some variability for particular questions. Academic researchers rated Mentorship & Teaching the highest across mastery, frequency, and importance, reflecting the fact that faculty spend a significant amount of time mentoring students and teaching in a classroom setting. Faculty note that they need mastery in mentorship and teaching and that these skills are important — not simply that they regularly use them.

Interestingly, neuroscience professionals regardless of job sector rated ‘Management and Communication’ skills to be strongly required, very important, and frequently used. Graduate students reported significantly less use and importance of management and communication skills than all other job sectors (Figure 4).

### Self-reported salaries correlate with skill profiles

Lastly, we asked whether the importance of skills for a given career would correlate with self-reported yearly salaries (Figure 5). Neuroscience professionals reported a range of salaries, with Applied/Industry scientists reporting the highest salaries (median=$150,000/year) followed by Consultants (median=$100,000/year; Figure 5a). There was a positive correlation between Salary and the importance of Coding (*p* = 0.023, r = 0.227), Data Analysis (*p* = 0.047, *r* = 0.199), and Management & Communication (*p* = 0.043, *r* = 0.203) skills (Figure 5b). There was a negative but not statistically significant correlation between Salary and the importance of Mentorship & Teaching skills (*p* = 0.441, *r* = −0.078). Trends for the frequency and mastery questions were similar, but only Data Analysis was significant (data not shown).

**Figure 5.**
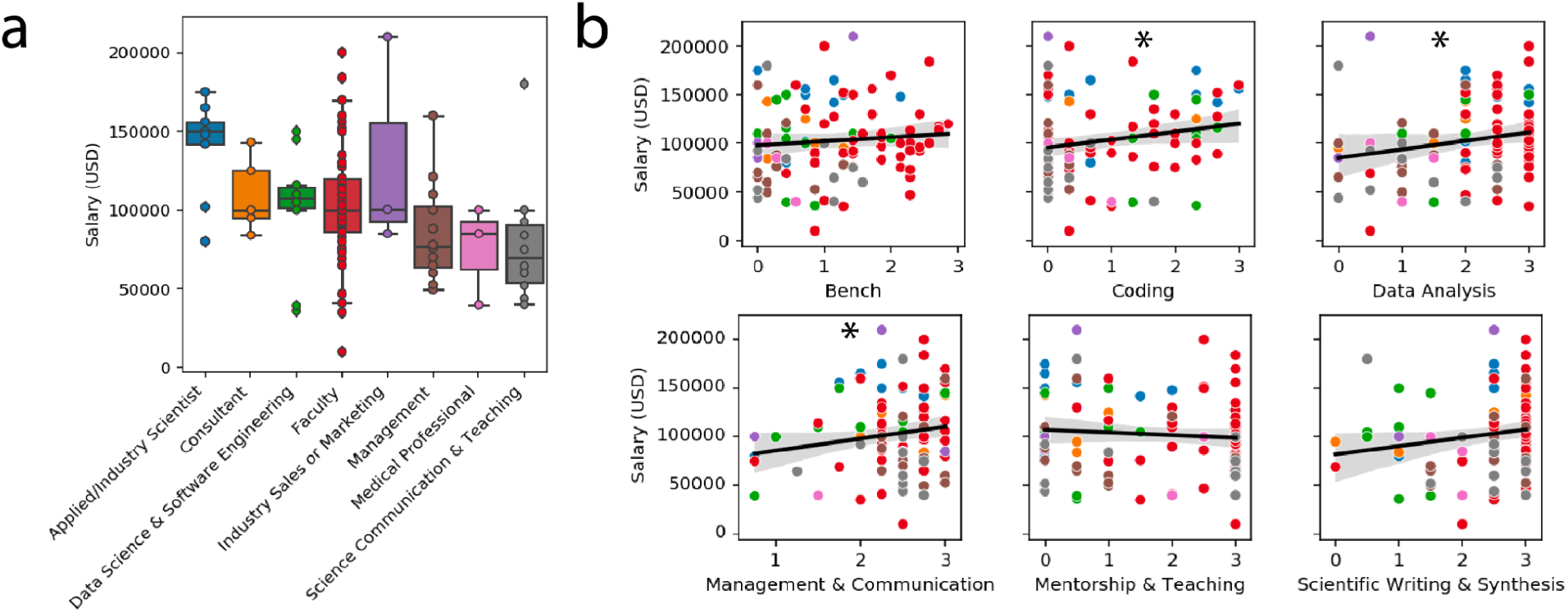
Relationship between yearly self-reported salaries (**a)** and importance of six different skill categories (**b**). Individual data points are colored by job category. Black line is the linear regression model fit, the gray shaded area is the 95% confidence interval of the fit. Asterisks indicate correlations at p<0.05 as tested by a Pearson correlation.

## Discussion

Here, we describe the skill profiles of hundreds of neuroscience graduate students and professionals in an effort to understand neuroscience career paths and inform our graduate training. We find that graduate students occupy very different skill spaces than neuroscience professionals, who tend to rely on more management and communication skills (Figures 1 & 5). While these differences may reflect the fact that more “soft skills” are required in the advanced stages of all careers, they also underscore the importance of these skills across career paths, and suggest that we should be training our graduates in these domains (Succi & Canovi, 2019). While others have noted the necessity of providing graduates with transferable skills such as being able to learn in groups and communication (e.g., Canelas et al., 2017; Scott et al., 2017, Watson & Burr, 2018), here, we provide evidence for this need specifically for careers of students with neuroscience degrees.

### Are we solely training graduate students for faculty careers?

In comparison to all of the neuroscience professionals, graduate students were most visibly similar to faculty members (Figures 1 & 5). This doesn’t come at much of a surprise, because graduate students are primarily trained by faculty, and graduate programs — particularly PhD programs — are almost exclusively designed to train future academic scientists. However, given that most PhD graduates will not end up in an academic job, the data we present here amplifies the call to provide more diverse training for our graduate students (Hoyne et al., 2016). Faculty also use coding, management, and communication skills, so integrating additional training in these domains would assist PhD students regardless of their career goals.

### Salary implications

Although faculty reported having, valuing, and using skills more than any other neuroscience professional, this is not reflected in most faculty salaries. Faculty salaries were highly variable, perhaps reflecting the vast differences in pay across universities (Johnson & Taylor, 2018). Rather, individual salaries could be predicted by the self-rated importance of three specific skill sets: Coding, Data Analysis, and Management and Communication. Coding and Data Analysis likely reflect the fact that jobs in this sector typically pay higher salaries, a trend which has been noted since the introduction of computers to the workplace (Krueger, 1993). Multiple job sectors in addition to Data Scientists & Software Engineers reported needing coding skills, which furthers an ongoing conversation about the need to teach coding to the next generation of scientists and knowledge workers (Akil et al., 2016; Grisham et al., 2016).

While the correlation between salary and Management & Communication skills may reflect the seniority of a professional moving up their career ladder, it may also echo previous observations that individuals with higher emotional intelligence earn higher salaries (Sanchez-Gomez et al., 2021). Regardless, it is important that trainees are aware of the salary implications associated with different skill sets and that all trainees have access to a wide array of professional development opportunities to improve on these skills.

### Improving graduate training

Given the observations here, there are several ways in which graduate training could be improved. First, students should be invited to develop “meta” work skills, intentionally working towards a skill set given their career goals (Bridgstock, 2009). The inclusion of practices such as the Individual Development Plan—when implemented well—are a good step in this direction (Vanderford et al., 2018; Tsai et al, 2018). Second, group work has been shown to enhance professional behaviors and job preparedness, particularly building communication and team management skills (Cartwright et al., 2021; Senay, 2015). Such group work could take place in the lab setting, as students work on research projects, or in the classroom.

Furthermore, providing graduate students with additional space to communicate their research, either verbally or in writing, is essential. Student-run writing groups such as NeuWrite (https://neuwrite.org/) or university-sponsored writing classes can give students necessary opportunities and critical feedback as they develop as writers. Finally, graduate students should be given access to and credit for coding classes; surprisingly, there are many PhD programs that do not offer any programming classes (SfN, 2017). It is also important to consider the timing of these skill interventions — students need to see the value in these skills in order to dedicate time and resources to learning them.

### Open questions

The work here raises several important questions that warrant further research. First, how do professionals learn each of these skills? While some of these skills may have been learned in the classroom or via professional development workshops, others may have been learned on the job. Understanding this would help us understand the accessibility of these skills, particularly for populations that have been historically excluded from STEM. Secondly, in this survey we did not ask about personal life skills such as time management or self-efficacy, which are clearly important in most STEM careers (Jang, 2016).

Perhaps most importantly, there are many open questions around the attitudes towards these skills, particularly in certain demographics. For example, a significant body of research has probed the perceptions of coding and computational career paths in Black and female students (Google Inc. & Gallup Inc., 20160; Baser, 2013; Cheryan et al., 2009). Given the clear salary implications, a better understanding of student perceptions not only has implications for the field of neuroscience, but for more broadly working towards a more equitable society.

## Acknowledgements

We would like to thank Dr. Qi Cui and the UC San Diego Chem/Bio Education Research Journal Club for useful early feedback on this manuscript, and Dr. Catherine Hicks for input on survey development. We would also like to thank the Faculty for Undergraduate Neuroscience (https://funfaculty.org/) and the neuroscience Twitter community for help disseminating this survey.

